# Integrated Identification of Disease Specific Pathways Using Multi-omics data

**DOI:** 10.1101/666065

**Authors:** Yingzhou Lu, Yi-Tan Chang, Eric P. Hoffman, Guoqiang Yu, David M. Herrington, Robert Clarke, Chiung-Ting Wu, Lulu Chen, Yue Wang

## Abstract

**Motivation:** Identification of biological pathways plays a central role in understanding both human health and diseases. Although much work has previously been done to explore the biological pathways by using single omics data, little effort has been reported using multi-omics data integration, mainly due to methodological and technological limitations. Compared to single omics data, multi-omics data will help identifying disease specific functional pathways with both higher sensitivity and specificity, thus gaining more comprehensive insights into the molecular architecture of disease processes.

**Results:** In this paper, we propose two computational approaches that integrate multi-omics data and identify disease-specific biological pathways with high sensitivity and specificity. Applying our methods to an experimental multi-omics data dataset on muscular dystrophy subtypes, we identified disease-specific pathways of high biological plausibility. The developed methodology will likely have a broad impact on improving the molecular characterization of many common diseases.

**Supplementary information:** Supplementary information attached.

## 1 Introduction

Pathway analysis is an important task for gaining novel insights into the molecular architecture of many complex diseases (Matthews, et al., 2009). With the advancement of new sequencing technologies, a large amount of quantitative gene expression data has been continuously acquired. The springing up omics data sets such as proteomics has facilitated the investigation of disease-relevant pathways.

Much work has previously been done by only exploring single omics data (Curtis, et al., 2005; Pollen, et al., 2014; Thurnherr, et al., 2016), but little has been invested into the study of interactions among multiple omics data, this is mainly due to the lack of computational power in the past to handle large-scale biological data and the complex biological relationship between molecules. While a single omics data can provide useful information about the underlying biological processes, multi-omics data integration would be much more comprehensive about the cause-effect processes responsible for diseases and their subtypes (Hasin, et al., 2017).

To find the disease-specific biological pathways with high sensitivity and specificity, this paper proposes two models to integrate miRNAseq, proteomics, and RNAseq data. These two models, the cascade model and the parallel model, are designed with different advantages and both can assess the statistical significance level of the identified pathways.

The cascade model improves both the ‘sensitivity’ and ‘specificity’. We first identify those pathways that are statistically significant in either mRNA(Shaw and Kamen, 1986) or protein (Washburn, et al., 2003), or significant in both sides, so sensitivity is improved. Next, in those selected pathways, we further take miRNA’s regulation into consideration, that is, only the pathways with some expected miRNA regulations (prior knowledge) would be kept, and this step helps improve specificity. On the other hand, our parallel model mainly focusses on improving ‘sensitivity’. In the parallel model, we explore the differentially expressed mRNAs/proteins, and mRNAs/proteins related to the miRNA regulations respectively. With each of the four sets of mRNAs/proteins, disease-specific pathways could be identified by gene set enrichment analysis. Then, a statistical method would be applied on these four sets of identified pathways to combine them, so that the sensitivity would be improved.

By applying our methods on the muscular dystrophy dataset, we successfully identified several pathways related to the disease. Hence, these two novel multi-omics data integration models and subsequent pathway identification will shed new light on pathophysiological processes in muscular dystrophies and improve our understanding on the molecular pathophysiology of muscle disorders, preventing and treating disease, and making people become healthier in the long term.

## 2 Relative Work

Several efforts have been made to explore pathways, but most pathways’ analysis was done solely on single omics data (Pollen, et al., 2014; Thurnherr, et al., 2016). This is mainly due to the high-level complexity within the regulatory relationship between these three molecules and the lack of technology to process a large amount of data in the past.

Thanks to technology improvement, it is capable and cost-efficient to obtain considerable data from microarray and RNAseq, which facilitates multi-omics data exploration. Since each kind of omics data would represent a reference and relationship with the disease, analyzing multi-omics can help us learn the causative change that induces the disease (Berger, et al., 2013; Hasin, et al., 2017), and would be more meaningful than solely reviewing single omics data.

Some efforts have been made to accumulate genomics data and explore omics data. In 2013, Berger et al. (Berger, et al., 2013) described some mathematics and statistical approaches for integrating genomics data. Bersanelli et al. (Bersanelli, et al., 2016) later classify diverse approaches of multi-omics integration into two categories. First is “network-based” theory which takes advantage of the currently known relationship to draw a graph to see the interactions between different variables. Second is the Bayesian model to calculate the posterior probability distribution for analyzing the multi-omics data(Bersanelli, et al., 2016). Even though these methods have considered the regulatory impact of miRNAs in the regulation of the pathway, they did not integrate both proteins and mRNAs, which are also disease specific for analyzing disease-related pathways. Hence, we proposed methods which have taken the complex regulation of mRNA, proteins, and miRNA as well as stringent mathematical models to find disease-specific mRNAs and proteins with the goal to detect disease-specific pathways.

## 3 Materials and methods

We proposed the two model: Cascade Model and Parallel Model to identify the pathways to integrate multi-omics data. Dataset are introduced the in 4.1 Experiment part.

### 3.1 Cascade Model

In the cascade model, we first perform the OVESEG test, a multiple-group comparison procedure, to define the disease-specific differential expressed proteins and mRNAs. Then, by performing gene set enrichment analysis on differentially expressed genes, we identify the pathways with mRNA or protein gene set enrichment analysis with a significant p-value. Next, we collect genes from those top pathways and further explore their relationship with miRNA. In the biological study, miRNA also plays a significant role in muscle development by regulating their target mRNAs and proteins. To exploit the relationship between miRNA and gene expression, we consult with the commonly used gene library – TargetScan (Lewis, et al., 2003) to collect all paired miRNA-mRNA and miRNA-protein co-expression pairs. Next, by conducting Pearson’s correlation coefficient, we measured the biologically expected correlation of each gene with its upstream miRNAs and identify those showing a negative correlation between the aforementioned miRNA-mRNA and miRNA-protein pairs. Furthermore, we identify and assess the most relevant disease-specific pathways by inputting negatively correlated genes into the gene-set libraries, and further characterize these prioritized marker subsets using KEGG (Kanehisa and Goto, 2000). We will then use Fisher’s method (Fisher, 2010) to combine pathways’ p-values derived from separate gene sets into a joint significance test assessing common pathway relevance. The cascade model would improve both the ‘sensitivity’ and ‘specificity’.

**Figure 1:**
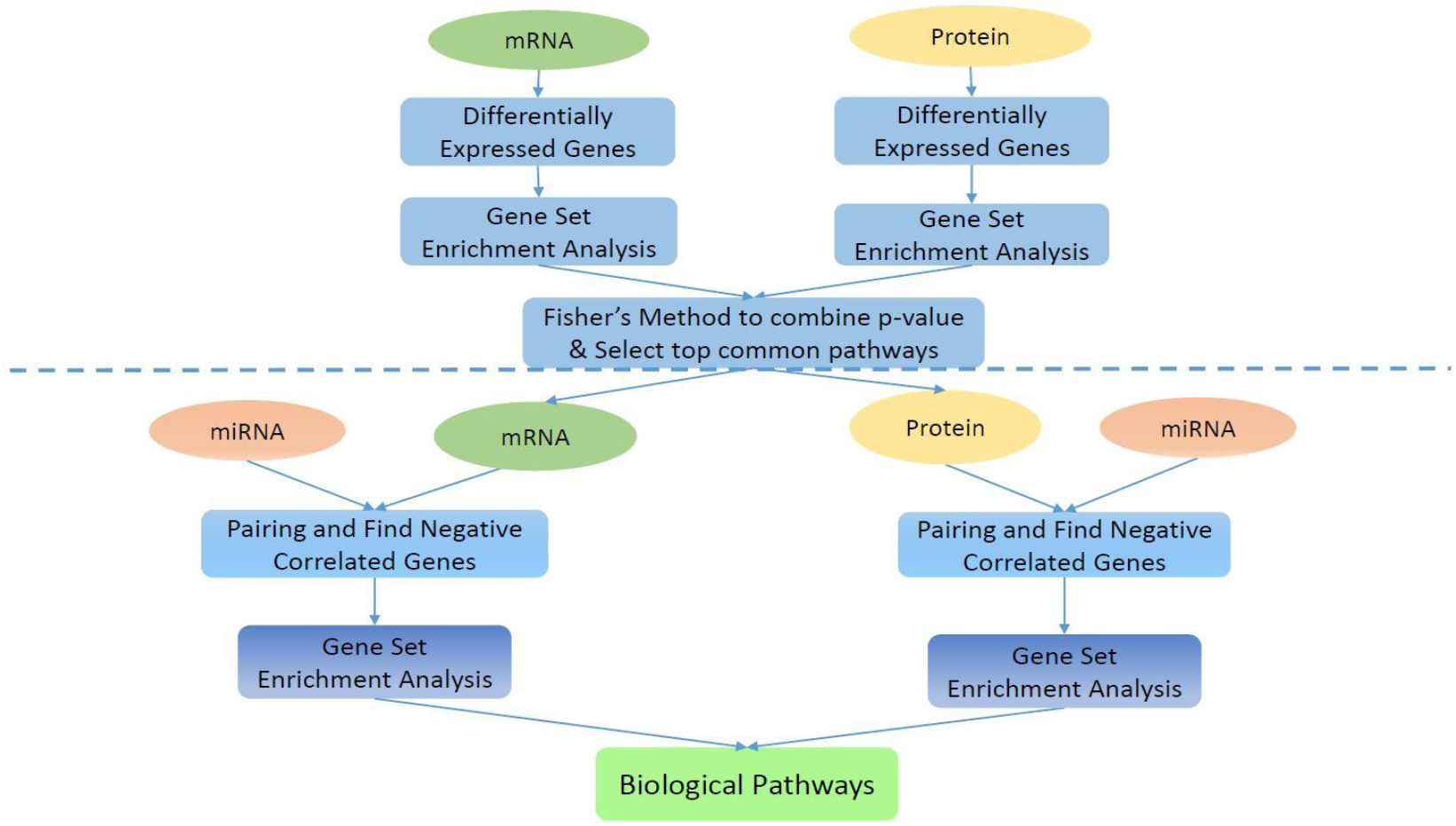
The workflow of multi-omics data integration-Cascade Model. The input are miRNA, mRNA and protein normalized data. We perform differential analysis on mRNA and protein, and conduct gene set enrichment analysis to find potential disease-specific pathways. Then we use Fisher’s method to combine p-values and select top pathways with the smallest p-values. For genes in these pathways, we find all the pairs of miRNA-mRNA and miRNA-protein from prior knowledge from TargetScan and calculate the correlation for each pair. After that, we pick up the mRNAs and proteins which are negatively correlated with their upstream miRNA respectively. Ultimately, we can identify biological pathways by gene set enrichment analysis.

### 3.2 Parallel Model

The parallel model mainly focusses on improving ‘sensitivity’. In the parallel model, OVESEG was performed to identify the differentially expressed mRNAs and proteins. Since miRNA also plays a significant role in the molecular study, to exploit the relationship between miRNA and gene expression, we combine prior knowledge to collect all miRNA with its target mRNA and protein. Next, we calculate the correlation coefficient for each upstream miRNA and its target mRNA/protein, and identify those mRNA/protein that shows a negative correlation in these pairs. Then after we identified the differentially expressed genes as well as genes that are biologically expected, we did the gene set enrichment analysis to find the pathways, and Fisher’s method was applied to have a joint p-value for each pathway from gene set enrichment analysis.

**Figure 2:**
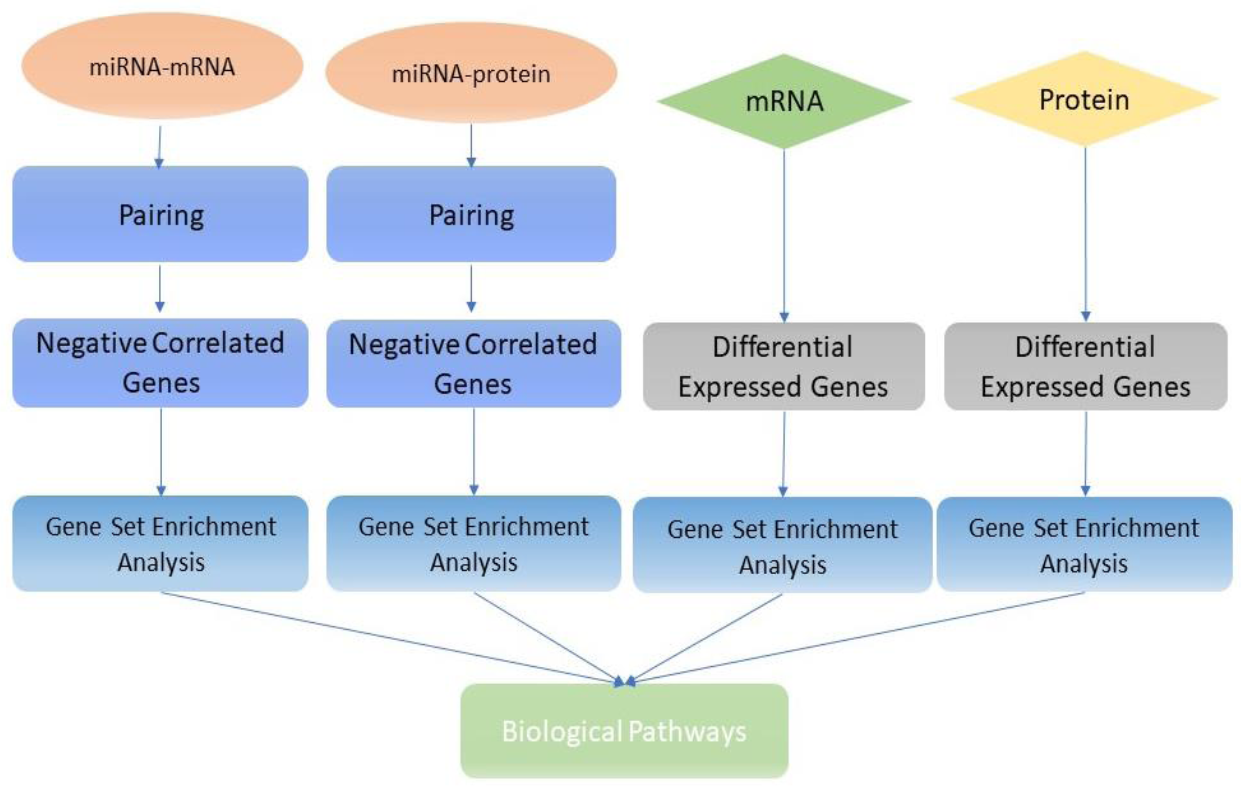
The workflow of multi-omics data integration – Parallel Model. The input are the same with cascade model. we performed differential analysis on mRNA and protein first, in addition, we find all the pairs of miRNA-mRNA and miRNA-protein from prior knowledge in the database - TargetScan. Then, we figure out the genes in mRNA and protein respectively negative correlated with its’ upstream miRNA. After that, we combine the selected genes from four separate lines and identify biological pathways by gene set enrichment analysis.

### 3.3 Differentially Expressed Genes

We use OVESEG (One Versus Everyone Subtype Exclusively-expressed Genes) to find disease-specific genes. The method is proposed by Chen (Lulu Chen, 2019), which used a novel permutation scheme to assess the statistical significance of marker genes (Lulu Chen, 2019; Yu, et al., 2011).

For each gene in mRNA and protein, the data are divided into three groups, the specific disease we investigate (like DMD or LMNA), normal, 6-other subtype diseases analysis. We are interested in the genes whose expression levels in a specific disease group significantly higher or lower than normal and other subtype disease groups. False discovery is likely to happen in multiple testing. To address this problem, the false discovery rate (FDR) suggested by Benjamini and Hochberg (Benjamini and Hochberg, 1995) are applied to this step. Here, *Q*-values can be generated by conducting multiple tests on p-values using false discovery rate control.

We conducted the differential analysis of each gene in mRNA and protein datasets, to ensure the genes we select are disease specific.

### 3.4 Correlation Analysis of Gene Pairs

Here we use Pearson’s correlation coefficient for selected the biologically expected genes. Pearson’s correlation coefficient (Pearson, 1895) can quantify both the direction and strength of the linear correlation between two variables. Since miRNA usually repress target gene expression, the negative correlation is biologically expected. We will find all mRNA and protein that has a negative correlation with its upstream miRNA.

### 3.5 Identification of negatively correlated mRNAs and proteins

In this step, we would like to identify those mRNAs and proteins which show a negative correlation with upstream miRNA, which is biologically expected. To identify correlation Pearson’s correlation coefficient (Pearson, 1895) are selected to quantify both the direction and strength of the linear correlation between two variables.

Referring to the TargetScan (Lewis, et al., 2003) database, matched microRNA and mRNA/protein profiling are obtained to visualize the miRNA and protein interactions. After knowing their interactions, we start analyzing their profiling under various conditions (Peter, 2010).There are three conditions in miRNA-mRNA / miRNA-protein pairing,

a. One-to-One miRNA-mRNA /protein pair
b. Multiple-to-One miRNA-mRNA/protein pair
c. One-to-Multiple miRNA-mRNA/protein pair

Condition a. is relatively easy to justify, by using fold change or correlation coefficient to see whether it is negatively correlated or not. Condition b. is more complicated. Bioinformatics analysis indicates a specific mRNA can be regulated by multiple miRNAs. We use the hypergeometric distribution to find the biological expected mRNAs and proteins. For condition c. because we only consider mRNA /protein genes as the unit of pathway enrichment analysis, this scenario is the same as a. after ungrouped.

In condition b., to identify significant genes which are negatively regulated by their corresponding miRNAs, we assume the test statistics to follow hypergeometric distribution under the null hypothesis,

For a given gene, the hypergeometric distribution is as followings

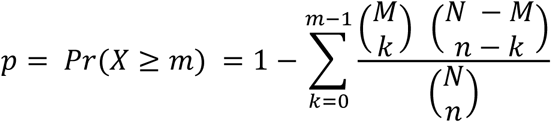

*N* = total number of miRNAs in the dataset

*M* = number of miRNAs negatively regulated with the mRNA among all the miRNAs in the dataset (without pairing)

*m* = number of miRNAs that negatively correlated with its’ targeted mRNA (with pairing)

*n* = number of miRNAs that targeted with mRNA

Each gene with its’ target miRNA pair will be given a p-value, small p-value means consider all the possible pairs, mRNAs or proteins are not negatively correlated to its upstream miRNA by chance, which is biologically expected.

### 3.6 Gene Set Enrichment Analysis

Gene set enrichment analysis is a commonly used method to identify known genes sets which are over-represented within a given sample of genes, which may have an association with disease phenotypes. The purpose of gene enrichment analysis is to retrieve the functional profile of that gene set with the goal of better understanding the underlying biological processes or molecular functions. Researchers use several statistical approaches and developed some software platforms to identify significantly enriched groups of genes. Typically, we use hypergeometric distribution for gene set enrichment analysis.

The probability of observing at least *x* genes in a particular pathway 

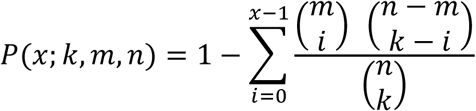

*n*: gene number in the database

*m*: gene number in the pathway

*k*: number of expected genes in the dataset (negatively correlated in miRNA-mRNA/ miRNA-protein)

*x*: genes numbers that are expected and also in the pathway

A pathway which has a p-value less than 0.05 would be selected as the statistically significant pathway. After research and comparison, we select Enrichr (Chen, et al., 2013) for gene set enrichment analysis. The gene lists are used as input to Enrichr for computing enrichment with prior knowledge from gene-set libraries. And the pathway database is the Kyoto Encyclopedia of Genes and Genomes (KEGG) (Kanehisa and Goto, 2000).

### 3.7 Fisher’s method to combine pathway p-values

Consider a set of *k* independent tests on the same null hypothesis. For each test, a significant p-value *p*_*i*_, is obtained(Fisher, 2010).

Fisher’s method combines p-values into one test statistics *χ*^2^ 

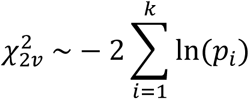

*χ*^2^ is a chi-squared distribution with 2*ν* degrees of freedom, assume *Z*_*1*_, *Z*_*2*_, …, *Z*_*ν*_, are independent and obey normal distribution, then *χ*^2^ equals to the sum of their squares 

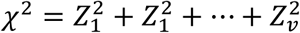

Where *ν*: degree of freedom, *ν* >, 0

The probability density function would be 

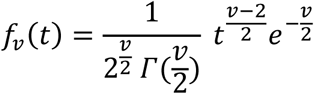

where *Γ*(·)is the Gamma Function computes chi-square Cumulative Distribution Function *P*(*χ*^2^|*ν*)at each of the values in *χ*^2^ using the corresponding degrees of freedom in *ν*

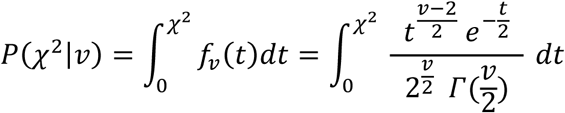

The upper tail *Q*(*χ*^2^|*ν*)

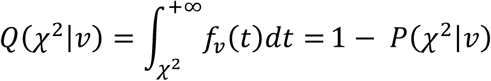

## 4 Experiment

Our experiment on the muscular dystrophy dataset corroborates the previous model we proposed. We are interested in finding out the molecular pathways that are related to each of the two diseases (DMD and LMNA) using the dataset we have in hand and identifying the pathways that are unique to each disease.

### 4.1 Datasets

The large-scale multi-omics dataset is derived from 40 human muscle biopsy tissue samples of 7 mutation-defined muscular dystrophies (DMD, COLVI, LMNA, RYR1, ANO5, BMD, CAPN3) and normal controls. Each biopsy has had generation of deep RNAseq-ribozero, microRNAseq, and SILAC proteomics (around 2,000 proteins quantitatively assayed in each biopsy), with a subset analyzed for mRNA splicing via long-range PacBio sequencing, and they carefully matched histopathology and choose 5 representative and well-preserved muscle biopsies per group.

Currently, our investigation is focused on two diseases, DMD and LMNA. Duchenne muscular dystrophy (DMD) is a type of dystrophin deficiency associated with muscle wasting with the voluntary muscles (Roberts, et al., 2015). The DMD disease type is the one we are interested in. DMD is a progressive muscular disease; at the same time, it is the most prevalent muscular dystrophy in children. The DMD biopsies were all in the early stage of disease (at diagnosis at about 5 years of age), and can be considered the most severe disease, thus showing the highest number of protein differences.

LMNA (lamin A/C) is another disease type we explored in multi-omics. LMNA causes Congenital Muscular Dystrophy (L-CMD) (Neuschwander-Tetri and Caldwell, 2003), which is a kind of genetic disorder displays in axial weakness, and may cause progressive muscle weakness and degeneration. In this paper, we will mainly show the result of DMD, while we attach the result of LMNA in the supplemental information part.

### 4.2 Identification of differentially expressed genes

The identification of differentially expressed genes is based on OVESEG. The method would find genes significantly higher or lower expressed in one disease group than normal or other disease groups. P-value and Z-score are calculated for each gene and adjusted for multiple testing corrections. When the p-value is very small (like smaller than 0.05), it represents the gene is significantly higher or lower in one group than another group, and it belongs to the expected differentially expressed genes. Q-values are computed from p-values with false discovery rate control, we set the threshold as 0.06. **Table 2** shows the genes differentially expressed in DMD.

**Table 1:**
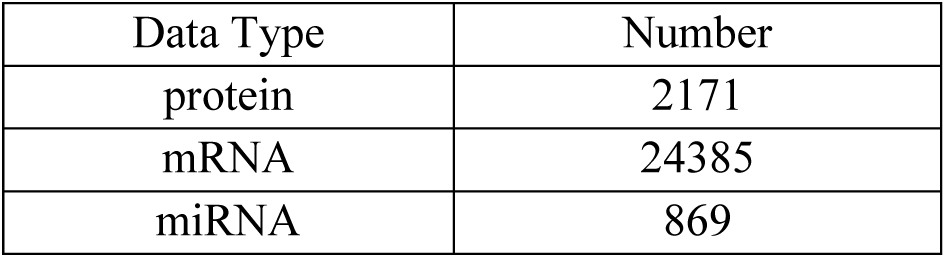
Dataset Composition

| Data Type | Number |
| --- | --- |
| protein | 2171 |
| mRNA | 24385 |
| miRNA | 869 |

**Table 2:** Differentially expressed mRNAs and proteins

|  | mRNA<br>(PUG) | mRNA<br>(PDG) | Protein<br>(PUG) | Protein<br>(PDG) |
| --- | --- | --- | --- | --- |
| Differentially Expressed Genes | 604 | 519 | 391 | 70 |

As **Table 2:** Differentially expressed mRNAs and proteins shows, after differential analysis and setting q-value 0.06 as the threshold, we can find 1123 mRNAs and 461 proteins as differentially expressed genes.

### 4.3 Cascade Model - Identification of pathways by differentially expressed genes gene set enrichment analysis

We conduct the gene set enrichment analysis for finding the potential pathways by inputting differentially expressed mRNAs and proteins separately to the database. We found 245 pathways enriched by mRNAs and 191 pathways enriched by proteins. There are 167 common pathways appeared both in differentially expressed mRNA and differentially expressed protein enriched pathways. In this step for a pathway, if its Fisher’s combined p-values smaller than 0.001 or p-value in the mRNA or protein’s gene set enrichment analysis smaller than 0.001, this pathway would be picked up. Using this criterion, 49 pathways are selected. Pathways numbers are listed in **Table 3**, detailed pathways are listed in supplemental information Table 3.

**Table 3:**
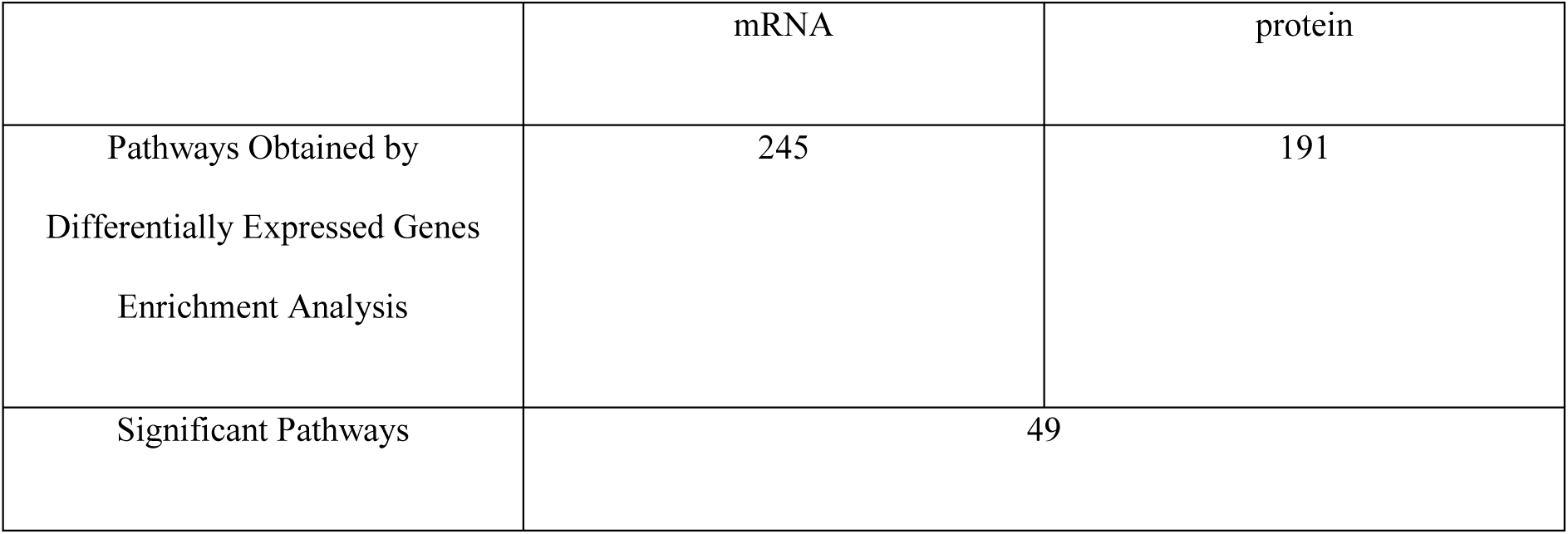
Pathways obtained by differentially expressed genes gene set enrichment analysis (database: KEGG)

**Table 4:**
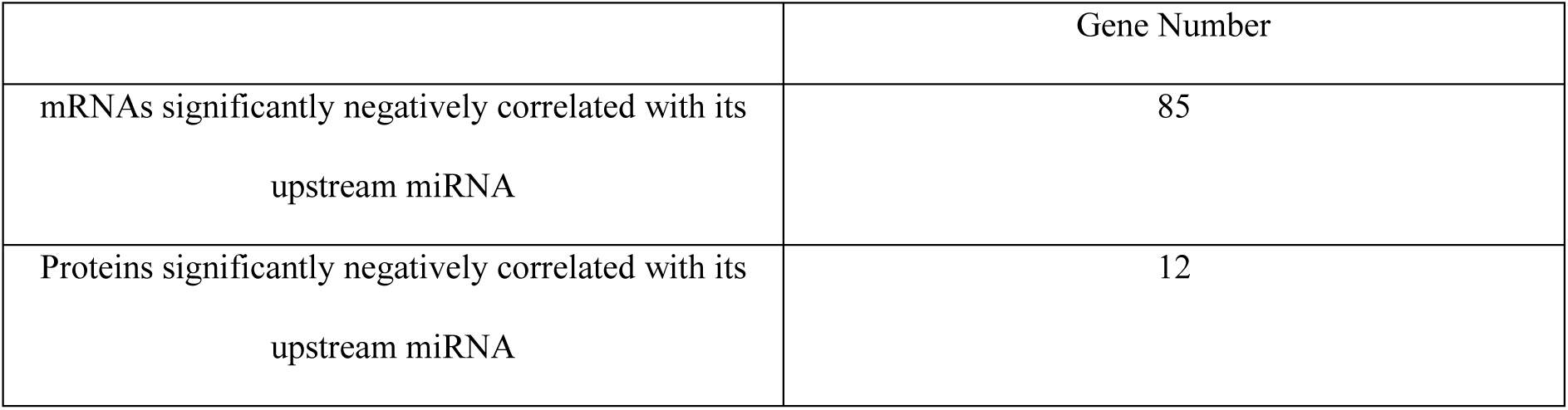
Number of genes negatively correlated with upstream miRNA

### 4.4 Cascade Model - Identification of significant negatively correlated genes

Within the 49 significant pathways from **Table 3**, we collect all the genes from those pathways refer to the KEGG pathways database. There are 2590 genes in total, from these genes, 2508 mRNAs and 380 proteins have expression data in our datasets.

We use selected mRNA and protein for conducting hypergeometric distribution for identifying gene in mRNA and protein which are negatively correlated with its upstream miRNA. In our experiment, there are 85 mRNAs, and 12 proteins significantly correlated with upstream miRNA (p-value<0.05).

The significant negatively correlated genes list with its p-value in mRNA and protein could be seen in supplemental material **Table 1** and **Table 2**.

### 4.5 Cascade Model - Upstream miRNA

miRNAs are a group of small noncoding RNAs (Bartel, 2009) that regulate their downstream target mRNA gene expression post-transcriptionally or proteins post-translationally. It is biologically expected that such miRNA-mRNA or miRNA-protein regulation is mainly suppression.

The muscular disease may be caused by multiple miRNAs cooperation function (Emery, 2002) identifying miRNAs that may also help us understand the mechanisms underlying specific disease. After we select the mRNAs and proteins respectively, we also retrieve their upstream miRNA for analysis. The number of upstream miRNAs for mRNA is 310 in total, and the upstream miRNA for proteins is 256 in total.

### 4.6 Cascade Model - Identification of pathways by negatively correlated genes gene set enrichment analysis

After we extract the genes from mRNA and protein which are negatively correlated with its’ upstream miRNA, we performed gene set enrichment analysis on those genes.

We found 166 pathways from significant negatively correlated mRNAs and 55 in proteins specific in DMD after gene set enrichment analysis.

Using Fisher’s method to combine the pathway obtained from significant mRNA gene set enrichment analysis and from significant mRNA gene set enrichment analysis, we can get the combined p-values for each pathway, and 33 significant pathways Fisher’s method combined p-value<0.001. Pathways list are attached in supplemental information **Table4**.

### 4.7 Cascade Model - Top Pathways

In **Table 6** we showed the top three pathways from differentially expressed genes and negatively correlated genes enrichment analysis. These pathways can be found in both differentially expressed genes enrichment analysis and negatively correlated genes enrichment analysis, with a significant combined p-value. The pathway list shows in supplemental information in **Table 3**. Supplemental information Table 5-8 lists the results for LMNA.

**Table 5:**
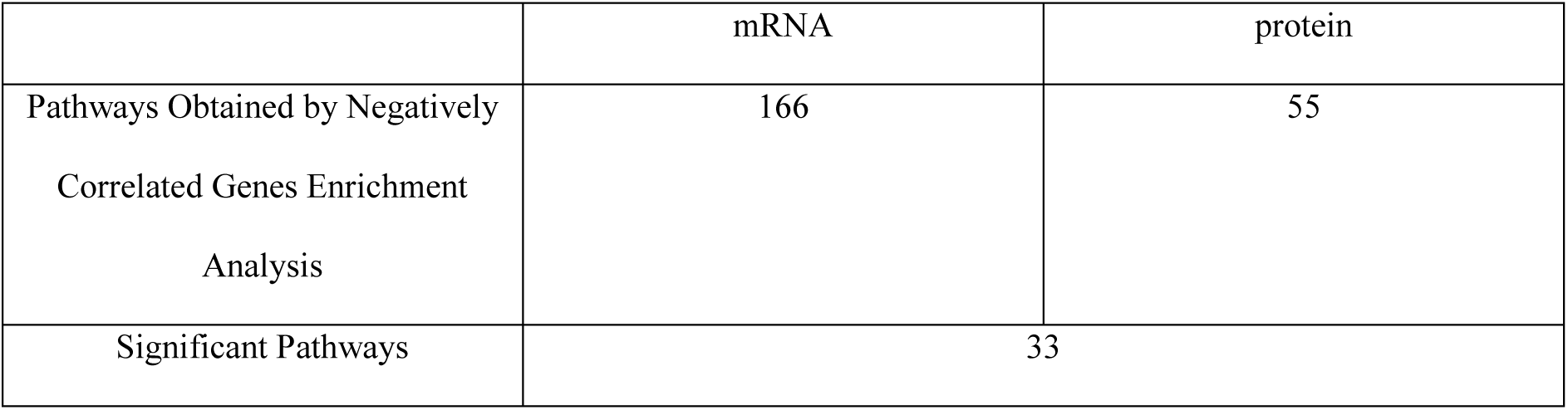
Pathways obtained by negatively correlated genes gene set enrichment analysis (database: KEGG)

|  | mRNA | protein |
| --- | --- | --- |
| Pathways Obtained by Negatively Correlated Genes Enrichment Analysis | 166 | 55 |
| Significant Pathways | 33 |  |

**Table 6:** Top pathways obtained by negatively correlated genes gene set enrichment analysis (database: KEGG)

|  | p-value from negatively correlated mRNA enrichment analysis get pathways | p-value from negatively correlated protein enrichment analysis get pathways | Fisher's method combined p-value |
| --- | --- | --- | --- |
| Calcium signaling pathway_Homo sapiens_hsa04020 | 8.17174E-03 | 2.40955E-09 | 5.05E-10 |
| Focal adhesion_Homo sapiens_hsa04510 | 1.33292073E-01 | 4.65066E-09 | 1.38E-08 |
| Pathways in cancer_Homo sapiens_hsa05200 | 2.2196475E-01 | 6.32576E-09 | 3.00E-08 |

**Table 7:** Common pathways specific in DMD

|  | Gene Number |
| --- | --- |
| mRNAs significantly negatively correlated with its upstream miRNA | 124 |
| Proteins significantly negatively correlated with its upstream miRNA | 25 |

**Table 8:** Top pathways obtained by negatively correlated genes gene set enrichment analysis might be related to muscular disease. (database: KEGG)

| Pathway Name | P-value<br>(pathway from mRNA gene set<br>enrichment analysis) | P-value<br>(pathway from protein gene set<br>enrichment analysis) | Combined P-value |
| --- | --- | --- | --- |
| Striated Muscle<br>Contraction_Homo<br>sapiens_WP383 | 0.225 | 0.050 | 0.062 |
| Amyotrophic lateral<br>sclerosis (ALS) | 0.211 | 0.046 | 0.055 |

### 4.8 Parallel Model - I Identification of significant negatively correlated genes

### 4.9 Parallel Model - Top Pathways

We found 24 common pathways from significant and negative correlated in mRNA and protein specific in DMD after gene set enrichment analysis. Here are the pathways which might be related to muscular disease. The pathways are listed in supplemental information Table 13.

## 5 Discussions

In this paper, we proposed two novel and systematic methods to identify disease-specific pathways by multi-omics data integration. Additionally, we explored the capacity of our model to identify the biological pathways from the muscular dystrophy data. By applying our method on the muscular dystrophy dataset, it is possible to find disease-related pathways by integrating multi-omics data. This study is the first step of large-scale data integration towards enhancing the understanding of the molecular pathophysiology of the more common muscle disorders. Later, we will validate our findings from biological studies.

Further research could also be undertaken in areas such as exploration of the mir-29 family. The mir-29 family has been found targeting a large group of functionally related genes, which effect on kidney, muscle and other organs. There are three members in the miR-29 family: miR-29a, miR-29b, and miR-29c. MiR-29b is the one that results in multiple types of muscle atrophy. Our research can further be applied to miR-29b (Li, et al., 2017). Furthermore, in the current models, we focus on the direct regulation between each miRNA and its target mRNA/protein, in the future, we will also take indirect regulation into consideration. To sum up, our model can identify biological pathways on real data, which may help gain novel insights into the molecular architecture of many complex diseases.

## ACKNOWLEDGMENTS

This work was funded in part by the National Institutes of Health under Grants HL111362-05A1, HL133932, W81XWH-18-1-0723 (BC171885P1), and CA184902-01.

