## Supplemental Information for "Integrated Identification of Disease Specific Pathways Using Multi-omics data"

### Multi-omics Data Integration for Identifying Disease Specific Biological Pathways

[1]Department of Electrical and Computer Engineering, Virginia Polytechnic Institute and State University, Arlington, VA 22203, USA; [2]School of Pharmacy and Pharmaceutical Sciences, State University of New York, Binghamton, NY, 13902, USA; [3]Department of Internal Medicine, Wake Forest University, Winston-Salem, NC 27157, USA; [4]Lombardi Comprehensive Cancer Center, Georgetown University, Washington, DC 20057, USA

† Equal contribution

* To whom correspondence should be addressed.

### Outline

#### Differentially Expressed Genes

We use OVESEG (One Versus Everyone Subtype Exclusively-expressed Genes) to find disease-specific genes. The method is proposed by Chen(et al.,2019), which used a novel permutation scheme to assess the statistical significance of marker genes (Lulu Chen, 2019; Yu, et al., 2011). The null distribution of the OVESEG statistic is the likelihood that the expected expression level of one gene is the same in the top two or more disease groups. OVESEG computes one-versus-everyone statistic and estimates its null distribution by weighted permutation scheme. The null distribution of the OVESEG -statistic is the likelihood that the expected expression level of one gene is the same in the top two or more disease groups (Wang, et al., 2016).
$\bar{s}_k(j)$ is the geometric mean of expression levels for the $j$th molecule under phenotype $k$, and $C_k$ is the set of $N_k$ samples under phenotype $k$.

$$\bar{s}_k(j) = \left(\prod_{i\in C_k} s_{ij}\right)^{\frac{1}{N_k}}$$

We set a threshold $t_{j(K)}$ as PUG's number,

$$t_{jk} = \min_{l\neq k}\left\{\frac{\hat{\mu}_{jk} - \hat{\mu}_{jl}}{\hat{\sigma}_j\sqrt{\frac{1}{N_k}+\frac{1}{N_l}}}\right\} = \min_{l\neq k}\{t\text{-}stat_{k,l}(j)\}, k = 1, \dots, K$$

$$\mathbb{J}_{PUG} = \frac{\bar{s}_k(j)}{\sqrt{\prod_{l\neq \mathrm{k}} s_k(l)}} \geq \ t_{j(K)}$$

where $\hat{\sigma}_j{}^2$is the estimated genewise variance, $Nk$ is the sample number under phenotype $k$, $N_l$ is the sample number under phenotype $l$ and the arithmetic mean of $log\,\overline{s_{ıȷ}}(j)$ is $\hat{\mu}jk = \ log\,\overline{s_k}\ (j)$. $stat_{k,l}(j)$ is t-statistic between phenotype $k$ and $l$. We call $t_{jk}$ as OVE PUG-statistic associated with phenotype $k$.
For each gene in mRNA and protein, the data are divided into three groups, DMD, normal, 6-other subtype diseases analysis. We are interested in genes whose expression levels are significantly higher or lower in the specific disease group than other groups. False discovery is likely to happen in multiple testing. To address this problem, the false discovery rate (FDR) suggested by Benjamini and Hochberg (Benjamini and Hochberg, 1995)**Error! Reference source not found.** are applied to this step. $Q$-values can be generated by conducting multiple tests on p-values using false discovery rate control.
We conducted the differential analysis of each gene in mRNA and protein datasets, to ensure the genes we select are disease specific.

#### Correlation Analysis of Gene Pairs

Pearson's Correlation Coefficient (Pearson, 1895) can quantify both the direction and strength of the linear correlation between two variables.
Suppose we have $t$ samples (including case and control group) for each gene,

For each gene pair, $X$ represents paired gene expression data from miRNA and $Y$ represents gene expression data from mRNA/protein.

$X: x_{1,} x_{2,} x_{3,} x_{4,} x_{5,\ldots,} x_{t,}$ $\qquad$ $Y: y_{1,} y_{2,} y_{3,} y_{4,} y_{5,\ldots,} y_{t,}$

The Pearson's correlation coefficient for this gene sample

$$r = \frac{\sum_{i=1}^{t}(x_i - \bar{x})(y_i - \bar{y})}{\sqrt{\sum_{i=1}^{t}(x_i - \bar{x}^2)}\sqrt{\sum_{i=1}^{t}(y_i - \bar{y}^2)}}$$

$$where\ \bar{x} = \frac{1}{t}\sum_{i=1}^{t} x_i\ , \qquad \bar{y} = \frac{1}{t}\sum_{i=1}^{t} y_i$$

The null hypothesis $H_0$ is

$$r = 0$$

and the alternative hypothesis $H_1$ is

$$r \neq 0$$

if $r < 0$, the pair is negative correlated, which indicates the increased expression levels of the gene and decrease of its' targeted miRNA, if $r > 0$, it would be defined as positive correlated, which refers to miRNA and protein/mRNA increase or decrease simultaneously.

The p-value can be calculated by t-distribution

$$t = \frac{r\sqrt{n-2}}{\sqrt{1-r^2}}$$

$r$ is the correlation coefficient, and $n$ is the number of observation.
Usually, the null hypothesis is rejected if the p-value is smaller than 0.05, indicating that mRNA and its' paired upstream miRNA shows a significantly negative correlation. The p-value is $2 \times P(T > t)$ where $T$ follows a $t$ distribution with n – 2 degrees of freedom.

**Cascade Model**

Table 1: through significance test, mRNAs which are negatively correlated with its' upstream miRNA and its p-value in hypergeometric distribution, p-value less than or equal to 0.05 (DMD specific)

| gene name | p-value | gene name | p-value |
|---|---|---|---|
| CEBPB | 2.42E-09 | CLEC4M | 0.014086 |
| LAMP2 | 5.5E-08 | AMHR2 | 0.014706 |
| ACVR2B | 7.06E-07 | COL9A3 | 0.015178 |
| SYNPO | 1.5E-06 | SOS1 | 0.015954 |
| COL4A3 | 1.2E-05 | BMP3 | 0.017681 |
| NOS1 | 1.35E-05 | ABCA2 | 0.018043 |
| GRM1 | 1.9E-05 | SOS2 | 0.018325 |
| RPL36A | 2.63E-05 | CCR4 | 0.018963 |
| PPP1CB | 3.37E-05 | WNT11 | 0.019052 |
| RBL1 | 0.000107 | UBE2D3 | 0.02022 |
| H3F3A | 0.000179 | LIFR | 0.020722 |
| MRPL19 | 0.000199 | SERPINA5 | 0.023244 |
| ERBB4 | 0.000447 | VARS | 0.024344 |

| | | | |
|---|---|---|---|
| TICAM1 | 0.000457 | PPP3CB | 0.024448 |
| GRIN2A | 0.000567 | PIAS4 | 0.02513 |
| MITF | 0.000745 | PRKCE | 0.026021 |
| AGPAT3 | 0.000767 | PAK6 | 0.026268 |
| TARSL2 | 0.000774 | PARS2 | 0.028319 |
| YOD1 | 0.000881 | H3F3B | 0.030716 |
| MRPL35 | 0.001242 | ATP6V1G2 | 0.030945 |
| RAC1 | 0.002055 | SPTA1 | 0.031815 |
| PDGFRB | 0.00211 | CAPZA2 | 0.034922 |
| F11 | 0.002335 | FGB | 0.035648 |
| SH3GLB1 | 0.002634 | COL4A2 | 0.036578 |
| LAMA1 | 0.005401 | PLCB1 | 0.036685 |
| UBQLN2 | 0.006183 | MYH7B | 0.039341 |
| SH3GL2 | 0.006264 | CD59 | 0.039688 |
| RAB22A | 0.00648 | AVPR1B | 0.040225 |
| IGSF5 | 0.006606 | MARVELD3 | 0.041623 |
| KAT2B | 0.007159 | OSM | 0.041842 |
| RELT | 0.007325 | IGF1 | 0.042331 |
| ACE | 0.007522 | C4BPB | 0.042997 |
| CTNNA3 | 0.007648 | SAR1B | 0.043079 |
| PTK2B | 0.008098 | RPL3L | 0.04365 |
| UBE2D1 | 0.008784 | NOS2 | 0.044418 |
| NGF | 0.009033 | PLAT | 0.045329 |
| PPP2R2B | 0.009322 | RAB8A | 0.047248 |
| CAMK2G | 0.01037 | ROCK2 | 0.047669 |
| TOLLIP | 0.011337 | BMP8B | 0.047954 |
| GAB1 | 0.011432 | MS4A1 | 0.049169 |
| GNAQ | 0.011602 | C9 | 0.049491 |
| FCER1G | 0.011669 | H3F3C | 0.05 |
| RBMXL3 | 0.012555 | | |

Table 2: through significance test, proteins which are negatively correlated with its' upstream miRNA and its p-value in hypergeometric distribution for gene set enrichment analysis, p-value less than or equal to 0.05 (DMD specific)

| gene name | p-value | gene name | p-value |
|---|---|---|---|
| RPS13 | 0.001064 | RPS25 | 0.014728 |
| RDX | 0.001286 | VDAC3 | 0.015091 |
| ACP1 | 0.001303 | CAMK2B | 0.027048 |
| VDAC1 | 0.002099 | NSFL1C | 0.030713 |
| LAMA2 | 0.011241 | TUBA8 | 0.033455 |
| VTA1 | 0.013348 | QARS | 0.038831 |

Table 3: Pathways obtained by differentially expressed genes gene set enrichment analysis (database: KEGG) (DMD specific)

| Pathway Name | p-value from differentially expressed mRNA enrichment analysis get pathway | p-value from differentially expressed protein enrichment analysis get pathway | Fisher's method combined p-value |
|---|---|---|---|
| Ribosome_Homo sapiens_hsa03010 | 1 | 1.92E-28 | 0 |
| Systemic lupus erythematosus_Homo sapiens_hsa05322 | 1.46E-15 | 0.077118 | 4.22E-15 |
| Staphylococcus aureus infection_Homo sapiens_hsa05150 | 1.83E-13 | 1 | 5.55E-12 |
| Osteoclast differentiation_Homo sapiens_hsa04380 | 1.43E-11 | 1 | 3.72E-10 |
| Phagosome_Homo sapiens_hsa04145 | 1.55E-08 | 0.002005 | 7.84E-10 |
| Pathogenic Escherichia coli infection_Homo sapiens_hsa05130 | 1 | 1.65E-10 | 3.88E-09 |
| Viral carcinogenesis_Homo sapiens_hsa05203 | 3.3E-08 | 0.014277 | 1.06E-08 |
| Lysosome_Homo sapiens_hsa04142 | 3.3E-08 | 0.520851 | 3.24E-07 |
| Alcoholism_Homo sapiens_hsa05034 | 1.39E-07 | 0.399186 | 9.85E-07 |
| Chemokine signaling pathway_Homo sapiens_hsa04062 | 8.52E-08 | 0.851775 | 1.27E-06 |
| Leishmaniasis_Homo sapiens_hsa05140 | 1.18E-07 | 1 | 2.00E-06 |
| Leukocyte transendothelial migration_Homo sapiens_hsa04670 | 0.000157 | 0.00106 | 2.76E-06 |
| Complement and coagulation cascades_Homo sapiens_hsa04610 | 2.95E-07 | 0.734486 | 3.54E-06 |
| Antigen processing and presentation_Homo sapiens_hsa04612 | 0.000203 | 0.001207 | 3.98E-06 |
| Aminoacyl-tRNA biosynthesis_Homo sapiens_hsa00970 | 1 | 2.51E-07 | 4.07E-06 |

| | | | |
|---|---|---|---|
| Protein processing in endoplasmic reticulum_Homo sapiens_hsa04141 | 1 | 2.51E-07 | 4.07E-06 |
| Spliceosome_Homo sapiens_hsa03040 | 1 | 2.76E-07 | 4.44E-06 |
| Tuberculosis_Homo sapiens_hsa05152 | 5.05E-07 | 0.734486 | 5.86E-06 |
| Platelet activation_Homo sapiens_hsa04611 | 2.95E-06 | 0.296308 | 1.31E-05 |
| Bacterial invasion of epithelial cells_Homo sapiens_hsa05100 | 1 | 1.02E-06 | 1.51E-05 |
| Fc epsilon RI signaling pathway_Homo sapiens_hsa04664 | 2.13E-06 | 1 | 3.00E-05 |
| Fc gamma R-mediated phagocytosis_Homo sapiens_hsa04666 | 0.000231 | 0.014277 | 4.49E-05 |
| Regulation of actin cytoskeleton_Homo sapiens_hsa04810 | 1 | 5.57E-06 | 7.30E-05 |
| RNA transport_Homo sapiens_hsa03013 | 1 | 5.94E-06 | 7.74E-05 |
| Hematopoietic cell lineage_Homo sapiens_hsa04640 | 6.5E-06 | 1 | 8.41E-05 |
| Salmonella infection_Homo sapiens_hsa05132 | 1 | 1.75E-05 | 0.000209 |
| B cell receptor signaling pathway_Homo sapiens_hsa04662 | 2.88E-05 | 1 | 0.00033 |
| Focal adhesion_Homo sapiens_hsa04510 | 1 | 4.10E-05 | 0.000455 |
| Toll-like receptor signaling pathway_Homo sapiens_hsa04620 | 5.67E-05 | 1 | 0.000611 |
| TNF signaling pathway_Homo sapiens_hsa04668 | 8.31E-05 | 1 | 0.000864 |
| Herpes simplex infection_Homo sapiens_hsa05168 | 0.000157 | 0.598961 | 0.000965 |
| Rheumatoid arthritis_Homo sapiens_hsa05323 | 0.000184 | 0.561711 | 0.00105 |

| | | | |
|---|---|---|---|
| Toxoplasmosis_Homo sapiens_hsa05145 | 0.000157 | 0.93415 | 0.00144 |
| Intestinal immune network for IgA production_Homo sapiens_hsa04672 | 0.000157 | 1 | 0.001531 |
| Carbon metabolism_Homo sapiens_hsa01200 | 1 | 0.000191 | 0.001825 |
| Adherens junction_Homo sapiens_hsa04520 | 1 | 0.000191 | 0.001825 |
| Endocytosis_Homo sapiens_hsa04144 | 1 | 0.000191 | 0.001825 |
| Influenza A_Homo sapiens_hsa05164 | 0.000231 | 0.920358 | 0.002008 |
| Phospholipase D signaling pathway_Homo sapiens_hsa04072 | 0.000284 | 0.772467 | 0.002069 |
| Rap1 signaling pathway_Homo sapiens_hsa04015 | 0.000632 | 0.364257 | 0.002158 |
| Jak-STAT signaling pathway_Homo sapiens_hsa04630 | 0.000231 | 1 | 0.002163 |
| Tight junction_Homo sapiens_hsa04530 | 1 | 0.000232 | 0.002176 |
| Apoptosis_Homo sapiens_hsa04210 | 0.000683 | 0.399186 | 0.002511 |
| Shigellosis_Homo sapiens_hsa05131 | 1 | 0.000449 | 0.003907 |
| Chagas disease (American trypanosomiasis)_Homo sapiens_hsa05142 | 0.0006 | 0.812175 | 0.004201 |
| Cytokine-cytokine receptor interaction_Homo sapiens_hsa04060 | 0.000521 | 1 | 0.004457 |
| Pertussis_Homo sapiens_hsa05133 | 0.000656 | 0.875035 | 0.004859 |
| Vibrio cholerae infection_Homo sapiens_hsa05110 | 1 | 0.000639 | 0.005336 |
| Legionellosis_Homo sapiens_hsa05134 | 1 | 0.000986 | 0.00781 |

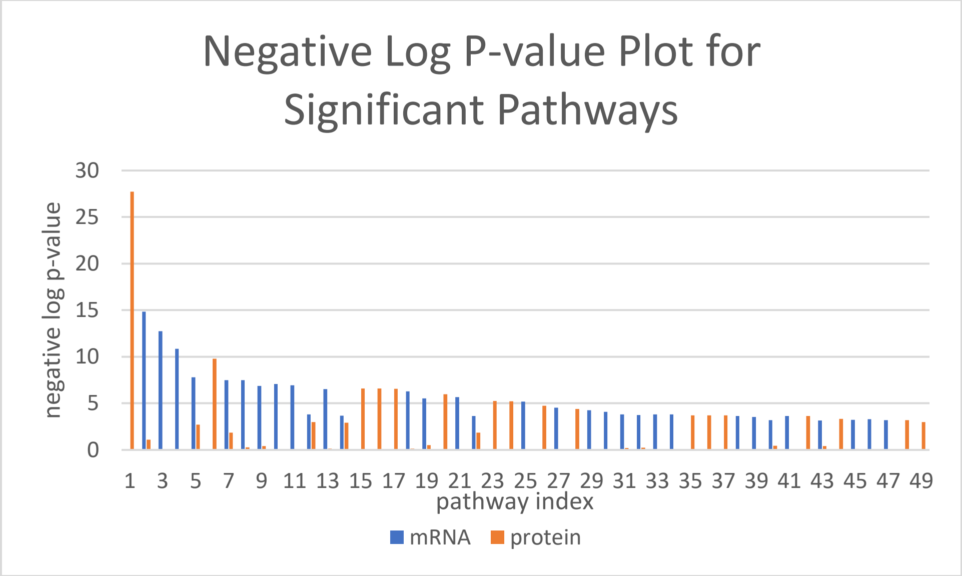

Figure 2 Plot of p-value after negative log 10 transform for significant pathways from differentially expressed genes gene set enrichment analysis

Table 4: Pathways obtained by negatively correlated gene set enrichment analysis (database: KEGG) (DMD specific)

| Term | p-value from negatively correlated mRNA enrichment analysis get pathway | p-value from negatively correlated protein enrichment analysis get pathway | Fisher's method combined p-value |
|---|---|---|---|
| Calcium signaling pathway_Homo sapiens_hsa04020 | 0.00817174 | 2.40955E-09 | 5.05E-10 |
| Focal adhesion_Homo sapiens_hsa04510 | 0.133292073 | 4.65066E-09 | 1.38E-08 |
| Pathways in cancer_Homo sapiens_hsa05200 | 0.22196475 | 6.32576E-09 | 3.00E-08 |
| Phospholipase D signaling pathway_Homo sapiens_hsa04072 | 1 | 2.23832E-08 | 4.17E-07 |
| Proteoglycans in cancer_Homo sapiens_hsa05205 | 0.05800596 | 4.15325E-07 | 4.47E-07 |
| Long-term potentiation_Homo sapiens_hsa04720 | 0.106620662 | 4.11153E-07 | 7.87E-07 |

| | | | |
|---|---|---|---|
| PI3K-Akt signaling pathway_Homo sapiens_hsa04151 | 0.19727012 | 5.09047E-07 | 1.72E-06 |
| Inflammatory mediator regulation of TRP channels_Homo sapiens_hsa04750 | 0.106620662 | 3.63739E-06 | 6.11E-06 |
| Amoebiasis_Homo sapiens_hsa05146 | 0.106620662 | 3.7621E-06 | 6.31E-06 |
| Complement and coagulation cascades_Homo sapiens_hsa04610 | 1 | 9.1492E-07 | 1.36E-05 |
| Dopaminergic synapse_Homo sapiens_hsa04728 | 0.114174272 | 1.63655E-05 | 2.65E-05 |
| ErbB signaling pathway_Homo sapiens_hsa04012 | 0.106620662 | 2.23665E-05 | 3.33E-05 |
| Gap junction_Homo sapiens_hsa04540 | 0.106620662 | 2.23665E-05 | 3.33E-05 |
| GnRH signaling pathway_Homo sapiens_hsa04912 | 0.106620662 | 2.55304E-05 | 3.76E-05 |
| Glioma_Homo sapiens_hsa05214 | 0.106620662 | 7.27085E-05 | 9.90E-05 |
| Ras signaling pathway_Homo sapiens_hsa04014 | 1 | 7.84348E-06 | 0.00010005 |
| Neurotrophin signaling pathway_Homo sapiens_hsa04722 | 0.11215782 | 8.45561E-05 | 0.000119171 |
| Tuberculosis_Homo sapiens_hsa05152 | 0.124362089 | 8.45561E-05 | 0.000131052 |
| Sphingolipid signaling pathway_Homo sapiens_hsa04071 | 1 | 1.08814E-05 | 0.000135239 |
| cGMP-PKG signaling pathway_Homo sapiens_hsa04022 | 0.05800596 | 0.000352458 | 0.000241202 |
| Wnt signaling pathway_Homo sapiens_hsa04310 | 0.117282048 | 0.000185378 | 0.000255165 |
| Regulation of actin cytoskeleton_Homo sapiens_hsa04810 | 0.135971417 | 0.000196079 | 0.000307464 |

| | | | |
|---|---|---|---|
| Circadian entrainment_Homo sapiens_hsa04713 | 0.106620662 | 0.000282913 | 0.000344141 |
| Melanogenesis_Homo sapiens_hsa04916 | 0.106620662 | 0.000320238 | 0.000385312 |
| Oxytocin signaling pathway_Homo sapiens_hsa04921 | 0.118919698 | 0.000299725 | 0.000400699 |
| Chagas disease (American trypanosomiasis)_Homo sapiens_hsa05142 | 1 | 5.15982E-05 | 0.000560977 |
| Long-term depression_Homo sapiens_hsa04730 | 1 | 5.15982E-05 | 0.000560977 |
| Renal cell carcinoma_Homo sapiens_hsa05211 | 1 | 7.44864E-05 | 0.000782472 |
| Amphetamine addiction_Homo sapiens_hsa05031 | 0.106620662 | 0.000750612 | 0.000834969 |
| Alcoholism_Homo sapiens_hsa05034 | 1 | 8.45561E-05 | 0.000877531 |
| Vascular smooth muscle contraction_Homo sapiens_hsa04270 | 1 | 8.45561E-05 | 0.000877531 |
| Platelet activation_Homo sapiens_hsa04611 | 1 | 8.92214E-05 | 0.000921156 |
| Chemokine signaling pathway_Homo sapiens_hsa04062 | 1 | 9.51198E-05 | 0.000975965 |

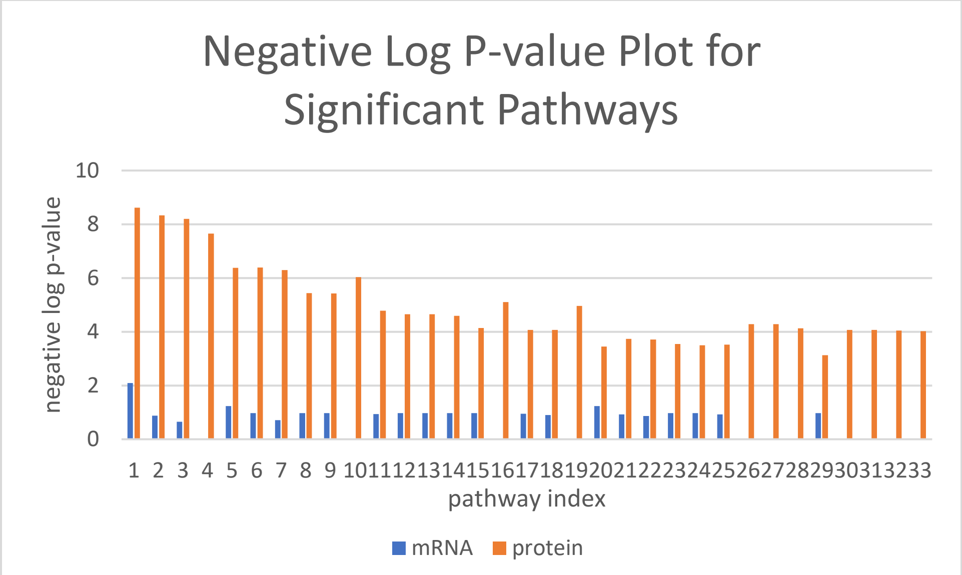

Figure 2 Plot of p-value after negative log 10 transform for significant pathways from negatively correlated genes gene set enrichment analysis

Table 5: through significance test, mRNAs which are negatively correlated with its' upstream miRNA and its p-value in hypergeometric distribution, p-value less than or equal to 0.05(LMNA specific)

| gene name | p-value | gene name | p-value |
|---|---|---|---|
| ATP6V1F | 1.49E-05 | WNT16 | 0.02116 |
| SDHA | 0.000311 | ACE | 0.022846 |
| GRIN2A | 0.000507 | PPP2R2B | 0.022986 |
| COX8C | 0.000737 | TPM4 | 0.02655 |
| FASLG | 0.001146 | FZD10 | 0.02712 |
| NOS1 | 0.001892 | COX10 | 0.028269 |
| ADRB1 | 0.002042 | WNT10A | 0.029251 |
| DNAI1 | 0.002184 | AKT3 | 0.038972 |
| TPM1 | 0.002231 | VDAC1 | 0.03925 |
| NRAS | 0.007343 | REST | 0.043179 |
| ADIPOR2 | 0.008649 | NDUFS2 | 0.045074 |
| CACNG4 | 0.010595 | WNT11 | 0.046185 |
| COX4I2 | 0.014033 | FRS2 | 0.049782 |

Table 6: through significance test, proteins which are negatively correlated with its' upstream miRNA and its p-value in hypergeometric distribution, p-value less than or equal to 0.05(LMNA specific)

| gene name | p-value | gene name | p-value |
|---|---|---|---|
| DMD | 3.29E-08 | DCTN2 | 0.012825 |
| ANK2 | 0.00013 | SGCA | 0.020165 |
| HSPG2 | 0.000867 | SGCD | 0.044092 |
| SGCB | 0.007511 | | |

Table 7: Pathways obtained by differentially expressed genes gene set enrichment analysis (database: KEGG) (LMNA specific)

| Pathway Name | p-value from differentially expressed mRNA enrichment analysis get pathway | p-value from differentially expressed protein enrichment analysis get pathway | Fisher's method combined p-value |
|---|---|---|---|
| Adrenergic signaling in cardiomyocytes_Homo sapiens_hsa04261 | 0.070376 | 0.095464 | 0.04033 |
| Parkinson's disease_Homo sapiens_hsa05012 | 0.070376 | 0.095464 | 0.04033 |
| Alzheimer's disease_Homo sapiens_hsa05010 | 0.191101 | 0.098251 | 0.093413 |
| Huntington's disease_Homo sapiens_hsa05016 | 0.205336 | 0.103071 | 0.102762 |
| Oxidative phosphorylation_Homo sapiens_hsa00190 | 0.264419 | 0.095464 | 0.118115 |
| Non-alcoholic fatty liver disease (NAFLD)_Homo sapiens_hsa04932 | 0.300281 | 0.095464 | 0.130489 |
| Metabolic pathways_Homo sapiens_hsa01100 | 0.264419 | 0.12402 | 0.144865 |
| Proteoglycans in cancer_Homo sapiens_hsa05205 | 0.401586 | 0.103071 | 0.173212 |

Table 8: Pathways obtained by negatively correlated genes gene set enrichment analysis (database: KEGG) (LMNA specific)

| Pathway Name | p-value from negatively correlated mRNA enrichment analysis get pathway | p-value from negatively correlated protein enrichment analysis get pathway | Fisher's method combined p-value |
|---|---|---|---|
| Arrhythmogenic right ventricular cardiomyopathy (ARVC)_Homo sapiens_hsa05412 | 0.11807 | 2.70E-08 | 6.56E-08 |

| | | | |
|---|---|---|---|
| Hypertrophic cardiomyopathy (HCM)_Homo sapiens_hsa05410 | 3.4E-05 | 2.87E-08 | 2.76E-11 |
| Dilated cardiomyopathy_Homo sapiens_hsa05414 | 4.4E-05 | 2.99E-08 | 3.69E-11 |
| Proteoglycans in cancer_Homo sapiens_hsa05205 | 1.7E-08 | 0.003747 | 1.57E-09 |
| Hepatitis B_Homo sapiens_hsa05161 | 0.00348 | 0.056252 | 0.001866 |
| Huntington's disease_Homo sapiens_hsa05016 | 1.3E-07 | 0.065635 | 1.66E-07 |

**Parallel Model**

Table 9: through significance test, genes in mRNAs which are negatively correlated with its' upstream miRNA and its p-value in hypergeometric distribution, p-value less than or equal to 0.05

| gene name | p-value | gene name | p-value |
|---|---|---|---|
| ABTB2 | 0.003011 | MTMR10 | 0.000114 |
| ACACB | 0.001586 | MUC3A | 0.04712 |
| ACSM2B | 0.038452 | NCKAP1 | 0.031804 |
| AGTPBP1 | 0.001676 | NDUFA4 | 0.025909 |
| AKAP7 | 8.67E-05 | NEK10 | 0.012225 |
| AMHR2 | 0.038833 | NMUR2 | 0.001794 |
| ANKRD40 | 0.049805 | NOS1 | 1.25E-05 |
| APOO | 0.008448 | NRG4 | 0.029836 |
| AQP4 | 0.001644 | NUDT8 | 0.013124 |
| ARL5A | 0.002728 | OTX1 | 0.010755 |
| ARNT2 | 4.58E-05 | PAQR9 | 0.004301 |
| ASB14 | 0.009951 | PGPEP1L | 6.51E-05 |
| ATP1B1 | 0.003783 | PHKB | 0.010419 |
| B3GALT1 | 0.00226 | PHTF2 | 0.035364 |
| B4GALNT1 | 0.002441 | PLCB1 | 0.0223 |
| BHLHE41 | 0.014581 | PLCL1 | 0.035159 |
| C11orf53 | 0.000945 | PLN | 0.000841 |
| C6orf10 | 0.013085 | PPARGC1A | 0.0414 |
| CAB39 | 0.000168 | PPP1CB | 4.07E-05 |
| CCDC91 | 0.000696 | PPP1R3A | 0.029633 |
| CLCN4 | 0.011072 | PPP6C | 0.014032 |
| CLIP4 | 0.000601 | PROP1 | 0.004052 |

| | | | |
|---|---|---|---|
| COX5A | 0.037394 | PTGES3 | 3.83E-05 |
| COX7C | 0.00895 | PTPN3 | 2.77E-05 |
| DCAF6 | 0.014018 | RCAN2 | 0.000457 |
| DHCR24 | 0.005393 | RCHY1 | 0.035264 |
| DMD | 0.037131 | REEP2 | 0.031776 |
| EIF3J | 0.012668 | RHOQ | 0.000313 |
| ENPP4 | 0.000103 | RIMS1 | 0.000195 |
| EPM2A | 0.001019 | RLN1 | 0.046605 |
| ERBB4 | 0.001908 | RMST | 0.009884 |
| FAM160A1 | 1.05E-08 | RNF128 | 4.6E-06 |
| FAM46C | 0.017061 | RNLS | 0.006529 |
| FBXO40 | 0.000149 | SAR1B | 0.044056 |
| FHOD1 | 0.024934 | SCN1B | 0.019981 |
| FKBP5 | 0.003636 | SCOC | 0.000249 |
| FOXP2 | 0.016959 | SDHD | 0.02031 |
| GAMT | 0.033559 | SEMA6B | 0.02716 |
| GAS2 | 7.44E-05 | SH3GL2 | 0.016481 |
| GBE1 | 0.012498 | SLC16A12 | 0.033505 |
| GOLGA7B | 1.94E-06 | SLC26A3 | 8.64E-06 |
| GOT2 | 0.048827 | SLC47A1 | 0.000471 |
| GTF2IRD2B | 0.004766 | SLITRK1 | 0.00652 |
| GUF1 | 4.19E-05 | SMOC1 | 0.000488 |
| HEMGN | 0.000239 | STAR | 0.008192 |
| INSM2 | 0.001472 | SYNPO | 5.45E-07 |
| KCNN2 | 0.003888 | TAOK2 | 0.032995 |
| KLHL32 | 0.000149 | TAPT1 | 0.033515 |
| KPNA5 | 0.03832 | TARSL2 | 6.29E-05 |
| LGR5 | 0.022633 | TDH | 0.003131 |
| LMO1 | 0.00022 | TEX2 | 0.00087 |
| LOC646736 | 0.031457 | THRB | 0.005166 |
| LYRM5 | 2.47E-05 | TMEM184A | 0.028079 |
| MAP3K9 | 9.54E-07 | TMEM201 | 0.008742 |
| MICAL1 | 0.04186 | TYRP1 | 0.008232 |
| MID2 | 0.00852 | UBE2D1 | 0.002822 |
| MITF | 0.00869 | UBE2Q2P2 | 0.015567 |
| MMADHC | 0.008921 | UHMK1 | 0.002583 |
| MN1 | 0.011415 | URM1 | 0.011478 |
| MOV10 | 0.039418 | USP13 | 0.002851 |
| MRPS30 | 0.019555 | XK | 0.00378 |

Table 10: through significance test, genes in proteins which are negatively correlated with its' upstream miRNA and its p-value in hypergeometric distribution, p-value less than or equal to 0.05

| gene name | p-value | gene name | p-value |
|---|---|---|---|
| ACP1 | 0.001303 | NDUFA10 | 0.002786 |
| CHCHD3 | 0.007979 | NDUFA6 | 0.040097 |

| | | | |
|---|---|---|---|
| CISD1 | 0.034334 | NDUFS8 | 0.027811 |
| COQ9 | 0.000253 | PDHA1 | 0.014596 |
| COX4I1 | 0.002095 | PDHX | 0.00574 |
| COX5A | 0.019495 | RPS13 | 0.001064 |
| DLST | 0.009934 | RPS25 | 0.014728 |
| DMD | 4.17E-07 | SGCD | 0.000449 |
| ESD | 0.01535 | SLC25A12 | 0.035356 |
| FTH1 | 0.006259 | SLC25A4 | 0.000148 |
| GOT1 | 0.00352 | SOD1 | 0.004876 |
| LMOD3 | 0.028721 | VDAC1 | 0.002099 |
| MTHFD1 | 0.018562 | | |

Table 11: Pathways obtained by negatively correlated mRNA gene set enrichment analysis (database: KEGG) (DMD specific)

| gene name | p-value | gene name | p-value |
|---|---|---|---|
| Ganglio Sphingolipid Metabolism_Homo sapiens_WP1423 | 0.002844 | Aryl Hydrocarbon Receptor Pathway_Homo sapiens_WP2873 | 0.249046 |
| Electron Transport Chain_Homo sapiens_WP111 | 0.00388 | Tryptophan metabolism_Homo sapiens_WP465 | 0.249046 |
| Farnesoid X Receptor Pathway_Homo sapiens_WP2879 | 0.006086 | Translation Factors_Mus musculus_WP307 | 0.249046 |
| Circadian rythm related genes_Homo sapiens_WP3594 | 0.008209 | Mecp2 and Associated Rett Syndrome_Mus musculus_WP2910 | 0.249046 |
| AMPK Signaling_Homo sapiens_WP1403 | 0.009052 | Myometrial Relaxation and Contraction Pathways_Homo sapiens_WP289 | 0.25208 |
| Glycogen Metabolism_Mus musculus_WP317 | 0.014809 | Diurnally Regulated Genes with Circadian Orthologs_Homo sapiens_WP410 | 0.258351 |
| Electron Transport Chain_Mus musculus_WP295 | 0.020147 | Diurnally Regulated Genes with Circadian Orthologs_Mus musculus_WP1268 | 0.258351 |
| Glycogen Metabolism_Homo sapiens_WP500 | 0.020938 | Exercise-induced Circadian Regulation_Mus musculus_WP544 | 0.258351 |
| Sulindac Metabolic Pathway_Homo sapiens_WP2542 | 0.030621 | Glycolysis and Gluconeogenesis_Mus musculus_WP157 | 0.258351 |
| ErbB signaling pathway_Mus musculus_WP1261 | 0.033049 | Mecp2 and Associated Rett Syndrome_Homo sapiens_WP3584 | 0.258351 |
| Arachidonate Epoxygenase / Epoxide Hydrolase_Homo sapiens_WP678 | 0.042607 | Glycolysis and Gluconeogenesis_Homo sapiens_WP534 | 0.26296 |
| ErbB Signaling Pathway_Homo sapiens_WP673 | 0.04575 | Synaptic Vesicle Pathway_Homo sapiens_WP2267 | 0.272094 |
| Effects of Nitric Oxide_Homo sapiens_WP1995 | 0.048545 | Translation Factors_Homo sapiens_WP107 | 0.272094 |
| FTO Obesity Variant Mechanism_Homo sapiens_WP3407 | 0.048545 | Oxidative phosphorylation_Mus musculus_WP1248 | 0.281115 |
| EV release from cardiac cells and their functional effects_Homo sapiens_WP3297 | 0.054447 | RANKL/RANK Signaling Pathway_Homo sapiens_WP2018 | 0.290025 |
| SREBF and miR33 in cholesterol and lipid homeostasis_Mus musculus_WP2084 | 0.060312 | Focal Adhesion-PI3K-Akt-mTOR-signaling pathway_Mus musculus_WP2841 | 0.290133 |

| | | | |
|---|---|---|---|
| Serotonin Transporter Activity_Homo sapiens_WP1455 | 0.066141 | Proteasome Degradation_Mus musculus_WP519 | 0.294439 |
| Alanine and aspartate metabolism_Homo sapiens_WP106 | 0.071934 | Kit receptor signaling pathway_Homo sapiens_WP304 | 0.307518 |
| Alanine and aspartate metabolism_Mus musculus_WP240 | 0.071934 | Endochondral Ossification_Mus musculus_WP1270 | 0.307518 |
| Alzheimers Disease_Mus musculus_WP2075 | 0.075402 | SIDS Susceptibility Pathways_Mus musculus_WP1266 | 0.307518 |
| SIDS Susceptibility Pathways_Homo sapiens_WP706 | 0.077519 | Oxidative phosphorylation_Homo sapiens_WP623 | 0.311824 |
| Insulin Signaling_Homo sapiens_WP481 | 0.077519 | Proteasome Degradation_Homo sapiens_WP183 | 0.320357 |
| Quercetin and Nf-kB/ AP-1 Induced Cell Apoptosis_Homo sapiens_WP2435 | 0.094752 | Endochondral Ossification_Homo sapiens_WP474 | 0.328785 |
| TCA Cycle_Homo sapiens_WP78 | 0.100369 | p53 signaling_Mus musculus_WP2902 | 0.328785 |
| SREBF and miR33 in cholesterol and lipid homeostasis_Homo sapiens_WP2011 | 0.105951 | Integrated Pancreatic Cancer Pathway_Homo sapiens_WP2377 | 0.334435 |
| Mitochondrial Gene Expression_Mus musculus_WP1263 | 0.105951 | TSH signaling pathway_Homo sapiens_WP2032 | 0.337109 |
| Amino Acid metabolism_Mus musculus_WP662 | 0.111496 | Rac1/Pak1/p38/MMP-2 pathway_Homo sapiens_WP3303 | 0.341232 |
| Urea cycle and metabolism of amino groups_Homo sapiens_WP497 | 0.117013 | Kit Receptor Signaling Pathway_Mus musculus_WP407 | 0.341232 |
| Mitochondrial Gene Expression_Homo sapiens_WP391 | 0.117013 | Primary Focal Segmental Glomerulosclerosis FSGS_Mus musculus_WP2573 | 0.34533 |
| Spinal Cord Injury_Mus musculus_WP2432 | 0.119582 | SREBP signalling_Homo sapiens_WP1982 | 0.357474 |
| Fatty Acid Biosynthesis_Homo sapiens_WP357 | 0.12794 | Primary Focal Segmental Glomerulosclerosis FSGS_Homo sapiens_WP2572 | 0.361471 |
| Fatty Acid Biosynthesis_Mus musculus_WP336 | 0.12794 | Leptin signaling pathway_Homo sapiens_WP2034 | 0.373318 |
| Transcription factor regulation in adipogenesis_Homo sapiens_WP3599 | 0.12794 | Arrhythmogenic Right Ventricular Cardiomyopathy_Homo sapiens_WP2118 | 0.373318 |
| XPodNet - protein-protein interactions in the podocyte expanded by STRING_Mus musculus_WP2309 | 0.129378 | G Protein Signaling Pathways_Mus musculus_WP232 | 0.422182 |
| Signal Transduction of S1P Receptor_Homo sapiens_WP26 | 0.144078 | Androgen receptor signaling pathway_Homo sapiens_WP138 | 0.42578 |
| Differentiation of white and brown adipocyte _Homo sapiens_WP2895 | 0.144078 | Corticotropin-releasing hormone_Homo sapiens_WP2355 | 0.436442 |
| Spinal Cord Injury_Homo sapiens_WP2431 | 0.168471 | G Protein Signaling Pathways_Homo sapiens_WP35 | 0.446908 |
| TCA Cycle_Mus musculus_WP434 | 0.17032 | Neural Crest Differentiation_Homo sapiens_WP2064 | 0.467264 |
| Alzheimers Disease_Homo sapiens_WP2059 | 0.17067 | Odorant GPCRs_Mus musculus_WP1397 | 0.532832 |
| p38 MAPK Signaling Pathway_Mus musculus_WP350 | 0.180591 | Adipogenesis genes_Mus musculus_WP447 | 0.550049 |
| Monoamine Transport_Homo sapiens_WP727 | 0.180591 | Adipogenesis_Homo sapiens_WP236 | 0.555647 |
| Constitutive Androstane Receptor Pathway_Homo sapiens_WP2875 | 0.180591 | Ectoderm Differentiation_Homo sapiens_WP2858 | 0.579995 |
| p38 MAPK Signaling Pathway_Homo sapiens_WP400 | 0.190737 | Regulation of Actin Cytoskeleton_Mus musculus_WP523 | 0.603019 |
| Endothelin Pathways_Homo sapiens_WP2197 | 0.190737 | Regulation of Actin Cytoskeleton_Homo sapiens_WP51 | 0.607963 |

| | | | |
|---|---|---|---|
| G13 Signaling Pathway_Mus musculus_WP298 | 0.195763 | Calcium Regulation in the Cardiac Cell_Mus musculus_WP553 | 0.607963 |
| Oxidative Damage_Mus musculus_WP1496 | 0.205722 | MAPK signaling pathway_Mus musculus_WP493 | 0.620057 |
| Nuclear Receptors_Mus musculus_WP509 | 0.205722 | EGF/EGFR Signaling Pathway_Homo sapiens_WP437 | 0.638642 |
| Amyotrophic lateral sclerosis (ALS)_Homo sapiens_WP2447 | 0.210656 | Chemokine signaling pathway_Mus musculus_WP2292 | 0.643146 |
| Nuclear Receptors_Homo sapiens_WP170 | 0.210656 | MAPK Signaling Pathway_Homo sapiens_WP382 | 0.649797 |
| Striated Muscle Contraction_Homo sapiens_WP383 | 0.210656 | EGFR1 Signaling Pathway_Mus musculus_WP572 | 0.656325 |
| G13 Signaling Pathway_Homo sapiens_WP524 | 0.220432 | TNF-alpha NF-kB Signaling Pathway_Mus musculus_WP246 | 0.67315 |
| BDNF signaling pathway_Homo sapiens_WP2380 | 0.224609 | mRNA processing_Mus musculus_WP310 | 0.710019 |
| Striated Muscle Contraction_Mus musculus_WP216 | 0.225275 | Non-odorant GPCRs_Mus musculus_WP1396 | 0.798589 |
| Tryptophan metabolism_Mus musculus_WP79 | 0.234872 | GPCRs, Class A Rhodopsin-like_Homo sapiens_WP455 | 0.802361 |
| Calcium Regulation in the Cardiac Cell_Homo sapiens_WP536 | 0.236033 | PluriNetWork_Mus musculus_WP1763 | 0.831182 |
| Myometrial Relaxation and Contraction Pathways_Mus musculus_WP385 | 0.240613 | PodNet: protein-protein interactions in the podocyte_Mus musculus_WP2310 | 0.852141 |
| Insulin Signaling_Mus musculus_WP65 | 0.247491 | | |

Table 12: Pathways obtained by negatively correlated protein gene set enrichment analysis (database: KEGG) (DMD specific)

| gene name | p-value | gene name | p-value |
|---|---|---|---|
| Electron Transport Chain_Mus musculus_WP295 | 1.42E-09 | TCA Cycle_Homo sapiens_WP78 | 0.021047 |
| Electron Transport Chain_Homo sapiens_WP111 | 2.63E-09 | Oxidative Stress_Mus musculus_WP412 | 0.032017 |
| Amino Acid metabolism_Mus musculus_WP662 | 4.93E-06 | One Carbon Metabolism_Homo sapiens_WP241 | 0.034439 |
| TCA Cycle_Mus musculus_WP434 | 6.85E-06 | One Carbon Metabolism_Mus musculus_WP435 | 0.034439 |
| Glycolysis and Gluconeogenesis_Mus musculus_WP157 | 2.88E-05 | Oxidative Stress_Homo sapiens_WP408 | 0.03806 |
| Glycolysis and Gluconeogenesis_Homo sapiens_WP534 | 3.06E-05 | Trans-sulfuration and one carbon metabolism_Homo sapiens_WP2525 | 0.039264 |
| Oxidative phosphorylation_Mus musculus_WP1248 | 3.88E-05 | Amyotrophic lateral sclerosis (ALS)_Homo sapiens_WP2447 | 0.04646 |
| Oxidative phosphorylation_Homo sapiens_WP623 | 5.63E-05 | Striated Muscle Contraction_Homo sapiens_WP383 | 0.04646 |
| TCA Cycle and PDHc_Homo sapiens_WP2453 | 0.000178 | Striated Muscle Contraction_Mus musculus_WP216 | 0.050038 |
| Folate Metabolism_Homo sapiens_WP176 | 0.003249 | One carbon metabolism and related pathways_Mus musculus_WP1770 | 0.058337 |
| Cytoplasmic Ribosomal Proteins_Mus musculus_WP163 | 0.003439 | Synaptic Vesicle Pathway_Homo sapiens_WP2267 | 0.061873 |
| Arrhythmogenic Right Ventricular Cardiomyopathy_Homo sapiens_WP2118 | 0.003936 | Vitamin B12 Metabolism_Homo sapiens_WP1533 | 0.064223 |
| Cytoplasmic Ribosomal Proteins_Homo sapiens_WP477 | 0.005496 | Copper homeostasis_Homo sapiens_WP3286 | 0.065396 |
| Acetylcholine Synthesis_Mus musculus_WP175 | 0.007478 | AGE/RAGE pathway_Homo sapiens_WP2324 | 0.07936 |

| | | | |
|---|---|---|---|
| Arachidonate Epoxygenase / Epoxide Hydrolase_Homo sapiens_WP678 | 0.008719 | mRNA processing_Mus musculus_WP310 | 0.087794 |
| Acetylcholine Synthesis_Homo sapiens_WP528 | 0.008719 | Selenium Micronutrient Network_Homo sapiens_WP15 | 0.10669 |
| Phase I biotransformations, non P450_Homo sapiens_WP136 | 0.009958 | Adipogenesis genes_Mus musculus_WP447 | 0.148377 |
| Trans-sulfuration pathway_Homo sapiens_WP2333 | 0.013668 | Adipogenesis_Homo sapiens_WP236 | 0.150518 |
| Iron Homeostasis_Mus musculus_WP1596 | 0.014901 | Ectoderm Differentiation_Homo sapiens_WP2858 | 0.160091 |
| Alanine and aspartate metabolism_Homo sapiens_WP106 | 0.014901 | NRF2 pathway_Homo sapiens_WP2884 | 0.167465 |
| Iron metabolism in placenta_Homo sapiens_WP2007 | 0.014901 | SIDS Susceptibility Pathways_Homo sapiens_WP706 | 0.182027 |
| Alanine and aspartate metabolism_Mus musculus_WP240 | 0.014901 | TNF-alpha NF-kB Signaling Pathway_Mus musculus_WP246 | 0.201398 |
| Dopamine metabolism_Homo sapiens_WP2436 | 0.016133 | XPodNet - protein-protein interactions in the podocyte expanded by STRING_Mus musculus_WP2309 | 0.267938 |

Table 13: Common Pathways with combined p-values

| Term | mRNA pathway P-value | Protein pathway P-value |
|---|---|---|
| Electron Transport Chain_Mus musculus_WP295 | 0.020147 | 1.42E-09 |
| Electron Transport Chain_Homo sapiens_WP111 | 0.00388 | 2.63E-09 |
| Amino Acid metabolism_Mus musculus_WP662 | 0.111496 | 4.93E-06 |
| TCA Cycle_Mus musculus_WP434 | 0.17032 | 6.85E-06 |
| Glycolysis and Gluconeogenesis_Mus musculus_WP157 | 0.258351 | 2.88E-05 |
| Glycolysis and Gluconeogenesis_Homo sapiens_WP534 | 0.26296 | 3.06E-05 |
| Oxidative phosphorylation_Mus musculus_WP1248 | 0.281115 | 3.88E-05 |
| Oxidative phosphorylation_Homo sapiens_WP623 | 0.311824 | 5.63E-05 |
| Arrhythmogenic Right Ventricular Cardiomyopathy_Homo sapiens_WP2118 | 0.373318 | 0.003936 |
| Arachidonate Epoxygenase / Epoxide Hydrolase_Homo sapiens_WP678 | 0.042607 | 0.008719 |
| Alanine and aspartate metabolism_Homo sapiens_WP106 | 0.071934 | 0.014901 |
| Alanine and aspartate metabolism_Mus musculus_WP240 | 0.071934 | 0.014901 |
| TCA Cycle_Homo sapiens_WP78 | 0.100369 | 0.021047 |
| Amyotrophic lateral sclerosis (ALS)_Homo sapiens_WP2447 | 0.210656 | 0.04646 |
| Striated Muscle Contraction_Homo sapiens_WP383 | 0.210656 | 0.04646 |
| Striated Muscle Contraction_Mus musculus_WP216 | 0.225275 | 0.050038 |
| Synaptic Vesicle Pathway_Homo sapiens_WP2267 | 0.272094 | 0.061873 |
| mRNA processing_Mus musculus_WP310 | 0.710019 | 0.087794 |
| Adipogenesis genes_Mus musculus_WP447 | 0.550049 | 0.148377 |

| | | |
|---|---|---|
| Adipogenesis_Homo sapiens_WP236 | 0.555647 | 0.150518 |
| Ectoderm Differentiation_Homo sapiens_WP2858 | 0.579995 | 0.160091 |
| SIDS Susceptibility Pathways_Homo sapiens_WP706 | 0.077519 | 0.182027 |
| TNF-alpha NF-kB Signaling Pathway_Mus musculus_WP246 | 0.67315 | 0.201398 |
| XPodNet - protein-protein interactions in the podocyte expanded by STRING_Mus musculus_WP2309 | 0.129378 | 0.267938 |

Lulu Chen, D.M.H., Robert Clarke, Guoqiang Yu,Yue Wang. Data-driven robust detection of tissue/cell-specific markers. *bioRxiv* 2019.

Pearson, K. Notes on regression and inheritance in the case of two parents. *Proceedings of the Royal Society of London* 1895;58 : 240–242.

Wang, N., *et al.* Mathematical modelling of transcriptional heterogeneity identifies novel markers and subpopulations in complex tissues. *Sci Rep* 2016;6:18909.

Yu, G., *et al.* PUGSVM: a caBIG analytical tool for multiclass gene selection and predictive classification. *Bioinformatics* 2011;27(5):736-738.
